# Visual representations in the human brain rely on a reference frame that is in between allocentric and retinocentric coordinates

**DOI:** 10.1101/2025.10.25.684527

**Authors:** Maria V. Servetnik, Michael J. Wolff, Chaipat Chunharas, Rosanne L. Rademaker

## Abstract

Visual information in our everyday environment is anchored to an allocentric reference frame – a tall building remains upright even when you tilt your head, which changes the projection of the building on your retina from a vertical to a diagonal orientation. Does retinotopic cortex represent visual information in an allocentric or retinocentric reference frame? Here, we investigate which reference frame the brain uses by dissociating allocentric and retinocentric reference frames via a head tilt manipulation combined with electroencephalography (EEG). Nineteen participants completed between 1728–2880 trials during which they briefly viewed (150 ms) and then remembered (1500 ms) a randomly oriented target grating. In interleaved blocks of trials, the participant’s head was either kept upright, or tilted by 45º using a custom rotating chinrest. The target orientation could be decoded throughout the trial (using both voltage and alpha-band signals) when training and testing within head-upright blocks, and within head-tilted blocks. Importantly, we directly addressed the question of reference frames via cross-generalized decoding: If target orientations are represented in a retinocentric reference frame, a decoder trained on head-upright trials would predict a 45º offset in decoded orientation when tested on head-tilted trials (after all, a vertical building becomes diagonal on the retina after head tilt). Conversely, if target representations are allocentric and anchored to the real world, no such offset should be observed. Our analyses reveal that from the earliest stages of perceptual processing all the way throughout the delay, orientations are represented in between an allocentric and retinocentric reference frame. These results align with previous findings from physiology studies in non-human primates, and are the first to demonstrate that the human brain does not rely on a purely allocentric or retinocentric reference frame when representing visual information.

## Introduction

Imagine visiting a big city and admiring its tallest building. As you gaze up, you may tilt your head to the side while taking in the impressive architecture. Unless you are looking at the tower of Pisa, the building stands upright relative to gravity. Put differently, when considered in an allocentric (or “world-centered”) reference frame, the building is oriented vertically. However, the information your visual system receives is the projection of the building onto your retinae. Incidentally, the sideways tilt of your head rotates this projection. As a result, in a retinocentric (or “retina-centered”) reference frame, the skyscraper is oriented diagonally. Despite the tilt of your head, you recognize that in allocentric coordinates the building is oriented vertically – retinocentric sensory input is transformed into an allocentric representation. Such transformation is necessary to maintain a continuous and stable experience of objects and scenes in the external world across a wide range of possible eye, head, and body movements. However, it remains largely unclear when and how this transformation from retinocentric to allocentric coordinates happens in the brain.

Human fMRI studies, which are performed with participants laying down and unable to move their heads or bodies, have tried to address the question whether visual information is represented in a retinocentric or allocentric reference frame by using eye movements to dissociate the two. For example, in an experiment by Golomb and Kanwisher^1^, participants were presented with a horizontal array of three possible stimulus locations, and they were told to fixate either to the left or to the right of the middle location (so either in between the two leftmost locations, or in between the two rightmost locations). The authors measured responses across visual cortex, including the primary visual cortex (V1), the intraparietal sulcus (IPS), and ventral object areas. In case of an allocentric representation that is anchored to the space external to the participant (also called “spatiotopic” in this kind of experiment), the response evoked by a stimulus shown in e.g., the leftmost possible location on the screen should be equivalent irrespective of where a participant is fixating. By contrast, in case of a retinocentric representation that stays anchored to the participant’s retinae in case of eye movements (also called “retinotopic” in this kind of experiment), the stimulus response should be equivalent across all possible stimulus locations on the screen, as long as the stimulus is at a fixed position (e.g., directly to the left) relative to where a participant is fixating. Results from this experiment^1^, and others like it^1–6^, largely suggest that visual stimuli are encoded and remembered in this latter, retinocentric reference frame, with limited evidence for allocentric coding (but see also^7,8^). However, people might rely more on an allocentric reference frame during tasks that more closely reflect the complexities of everyday naturalistic visual experience. For example, psychophysical work has shown that people remember locations more precisely in an allocentric compared to a retinocentric reference frame when multiple stimuli need to be remembered during an eye movement^9^, or when landmarks are present^10,11^. This suggests that, under certain circumstances, a transformation from a retinocentric to allocentric reference frame can be observed when eye movements are used to dissociate the two.

While eye movements might be a convenient way to tease apart retinocentric and allocentric reference frames, especially in experimental setups that greatly restrict other types of body movements (i.e., fMRI), eye movements may not be a representative motor manipulation to uncover reference frame transformations. It has been suggested that the reference frame for visual representations depends on the coordinates used by the end motor effectors^12^ (but see also^6^). For eye movements, these coordinates are likely retinocentric^13^, making a transformation unnecessary. This means that other types of body movement may be necessary to investigate reference frame transformations. Head movement is a particularly promising manipulation, given that the brain’s two main sources of information about head position are proprioceptive or corollary discharge signals from the neck muscles and vestibular system^14–19^, both of which are unlikely to operate in a retinotopic reference frame. Considering different types of body movements also means that, in addition to retinocentric and allocentric representations, we should acknowledge egocentric representations that are anchored to the body. The fact that different body parts can move independently of one another means that egocentric representations can be anchored to different body parts (e.g., can be body-centric, head-centric, effector-centric, etc.)^20–23^. For our purposes it’s important to note that when the head is the *only* body part that moves, and the eyes remain still inside of the head (i.e., move with it) the head-centric reference frame becomes equivalent to retinocentric reference frame (the retinae are anchored to the head). Similarly, the body-centric reference frame becomes equivalent to the allocentric reference frame (as the body is anchored to space).

Given that we need to understand our visual world across this range of possible reference frames, visual cortex is unlikely to represent visual inputs in an exclusively retinocentric way. Indeed, a growing body of monkey electrophysiology work shows that the parietal cortex processes proprioceptive and vestibular signals about head position^24,25^ and integrates these with visual signals^26^. Rosenberg and Angelaki^27^ demonstrated that, during a head-and-body tilt manipulation, the tuning functions of planar-tilt-selective neurons in the central intraparietal cortex (CIP) of monkeys would shift to account for a change in the direction of the gravity vector. However, rather than shifting completely over to an allocentric reference frame, these neurons maintain a heterogeneous set of representations, ranging from completely retinocentric to completely allocentric, with most tuning functions shifting partially.

There is parallel behavioral evidence from humans that retinocentric and allocentric reference frames are combined to inform visual perception and memory. When human participants were asked to encode a spatial position shown on a computer screen, and then underwent a head-and-body tilt manipulation (e.g., they were rotated 16º counterclockwise) before reporting the location relative to their head-and-body position, their reports were biased in the direction of their tilt^28^. Such a shift would not be expected if encoding happened in a purely retinocentric reference frame. Similar response shifts have been shown when only the head (and not the entire body) was tilted^29^. Thus, when retinotopic and spatiotopic reference frames are misaligned due to head or body tilts (instead of eye movements), visual processes no longer seem to be predominantly retinocentric, but reflect a combination of both retinocentric and allocentric reference frames. When these other body parts move, the pull of gravity could provide potentially stable and reliable signals to maintain an allocentric reference frame, which does not come into play when only the eyes are moved.

Relatedly, several studies have used the “oblique effect”^30,31^ to better understand human visual processing across different reference frames. The “oblique effect” refers to better discriminability for cardinal orientations (i.e., vertical and horizontal) compared to oblique orientations (i.e., diagonals). For example, Mikellidou et al.^32^ have shown that the overall sensitivity profile to different orientations shifts when independently manipulating head and body position, from which they concluded that visual information is represented in a combination of differentially weighted retinocentric and allocentric signals. In addition to higher precision around cardinals (the oblique effect), people also tend to bias their responses away from cardinals (the repulsion effect)^33,34^. Using a head tilt manipulation, Rademaker et al.^35^ showed that, while the oblique effect was anchored to a retinocentric reference frame (i.e., precision was highest for orientations that were cardinal on the retinae, irrespective of head tilt), the repulsion effect was anchored to an allocentric reference frame (i.e., responses were biased away from world-centered vertical and horizontal). This work implies that different stages of visual processing might rely on different reference frames. Specifically, it suggests that the oblique effect is more perceptual in nature – “hard-wired” into early visual processing over the course of development^36–38^, while the repulsion effect might be implemented at a later stage of visual processing where biases are referenced to the external world. Taken together, studies with head and body tilt manipulations in monkeys and humans imply that visual information is encoded in a combination of both retinocentric and allocentric coordinates.

Finally, what is largely overlooked in studies on visual reference frames is the distinction between visual information that is directly perceived, and visual information that is held in mind over a brief delay. In the simplest case, the reference frame(s) used during perception carry over into the delay. Alternatively, a reference frame transformation takes place once visual information is no longer tethered to perceptual input. There is increasing evidence that, from perception to memory, visual representations undergo a transformation^39–42^ and become less veridical and more categorical^43,44^. Might there be a similar transformation from perception to memory in terms of the reference frames used for visual information? It would make sense if the first feedforward sweep of information from the eyes is representing inputs in a retinocentric manner. Similarly, when there is no input from the eyes and visual information is held only in mind, might this information be coded in a more allocentric manner?

In the present study, we set out to examine which reference frame(s) may be used to encode visual information, and whether a possible transformation in reference frames occurs from perception to memory. To this end, we showed people grating stimuli of different orientations, and asked them to recall each orientation after a delay. During this task, we recorded their brain activity at high temporal resolution using electroencephalography (EEG). Critically, on half of the trails people tilted their head by 45º. Through the use of multivariate pattern analyses, we were able to access the reference frame in which visual orientations were encoded in the human brain from one moment to the next. We show, for the first time, that orientation information is coded in a combined allocentric and retinocentric reference frame in humans. Moreover, this combined reference frame is evident from the earliest time points of visual processing, and persists during the memory delay. These findings contradict previous neuroimaging studies that claim retinocentric coding of visual information via the use of eye movement manipulations. Instead, our findings are in line with monkey physiology and human behavioral studies using head-and-body tilt manipulations, and show that visual information is stably encoded in a combined allocentric and retinocentric reference frame throughout both perception and memory.

## Results

To assess whether visual information is represented in an allocentric or retinocentric reference frame in the human brain (Figure 1A), and to evaluate possible reference frame shifts between perception and working memory, we recorded electroencephalography (EEG) during a simple working memory task (Figure 1B). Participants viewed (150 ms), and then remembered (2000 ms), a randomly oriented target grating. At report, they rotated a recall dial to precisely match the orientation of this target. Importantly, our participants performed this task with their heads either upright or tilted at a 45º angle around the nasion, which allowed us to dissociate the allocentric from the retinocentric reference frame. Participants performed well on this task, and there were no significant differences in recall performance (*t*_(18)_ = -0.384, *p* = .705) or median reaction times (*t*_(18)_ = 0.378, *p* = .71) as a function of head tilt (Figure 1C).

**Figure 1.**
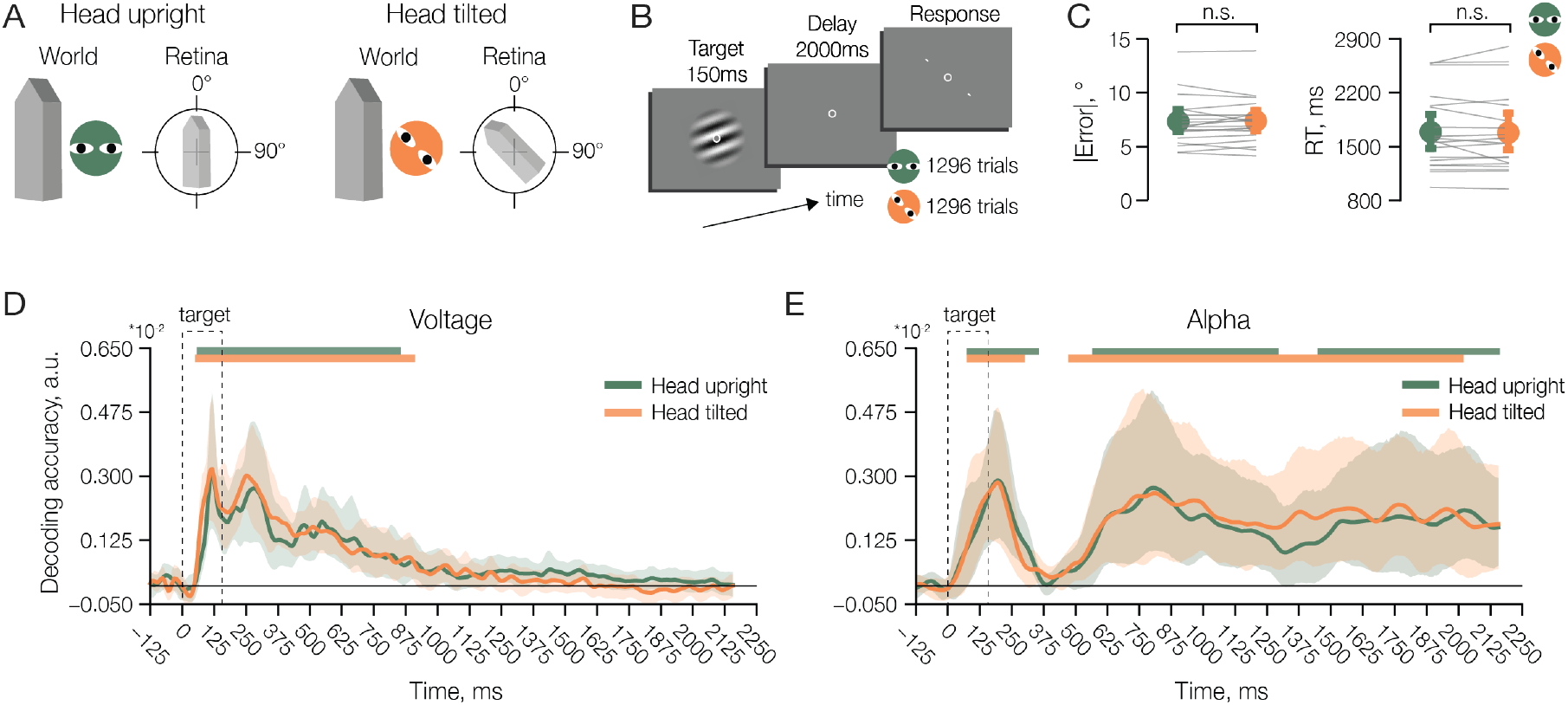
A: Visual items have an orientation in an allocentric reference frame – e.g., buildings are vertical relative to gravity. During visual perception, such items are projected on the retina, and as long as the head is upright, the orientation in the allocentric reference frame (e.g., vertical) matches the orientation in the retinocentric reference frame (e.g., vertical). However, when the head is tilted (by e.g., 45º), the projection on the retina rotates in the opposite direction of the tilt, resulting in a dissociation between the allocentric (e.g., vertical, or 0º) and retinocentric (e.g., -45º) reference frames. **B**: On every trial of the task, participants viewed a pseudo-randomly oriented target grating for 150 ms and then remembered this orientation over a delay of 2000 ms. Participants subsequently rotated a dial to match the target orientation. Importantly, participants performed half of the trials with their head upright, and half of the trails with their head tilted by 45 degrees (n = 9 tilted clockwise, n = 10 tilted counterclockwise). **C**: There were no significant differences in recall error (7.31º with 6.45º–8.41º CI, and 7.36º with 6.51º–8.31º CI) or response times (1688.7 s with 1453–1916.7 CI, and 1679.5 s with 1466.7–1923.4 CI) between the head-upright and head-tilted conditions, respectively. Gray lines indicate individual participants, while colored circles and error bars indicate across-participant averages +95% bootstrapped confidence intervals. **D**: Decoding accuracy over time for a decoder trained and tested on the EEG voltage signal. Both a decoder trained and tested on head-upright trials (green), and a decoder trained and tested on head-tilt trials (orange) reveal significant orientation information from about 50-1000 ms. after stimulus onset. **E**: Decoding accuracy over time (as in **D**), but for a decoder trained and tested on the alpha-band power. Here, the target orientation is briefly decodable during stimulus presentation (∼100-300 ms), and throughout most of (head upright) or the entire (head tilted) delay. For **D** and **E**, error bars/areas represent bootstrapped 95% confidence intervals of the mean, while green and orange bars denote significant decoding clusters for the head upright and head tilted trials, respectively (permutation test, n = 19, cluster-defining threshold *p* < .05, corrected significance level *p* < .05).

We confirmed that the target orientation was decodable from the EEG signal both during perception and memory. Specifically, we applied a pattern-similarity-based decoder, trained and tested within each head-tilt condition, to both the voltage (broadband) and the alpha-band (8-12 Hz) signals from the 17 posterior EEG channels (Figure 1D and 1E, respectively)^45^. With respect to the voltage signals, the target orientation was significantly decodable in both head-upright and head-tilted trials for approximately 1000 ms after stimulus onset (Figure 1D). This result is in line with previous studies showing that, while directly viewed stimuli are well decodable from EEG voltage signals, mnemonic information tends to decay^45,46^. This decoding was not driven by small eye movements (as implied by some previous work^47^; Supplementary Figure 1A). With respect to the alpha-band (8-12 Hz) signals, the target orientation was significantly decodable from early during the delay (∼500 ms after stimulus onset) until the end of the delay (2150 ms after stimulus onset) (Figure 1E). There was also an earlier increase in decoding (∼100-300 ms after stimulus onset), which likely reflects an “evoked” alpha oscillation from the stimulus presentation. This result is in line with previous research showing that alpha-band (8-12 Hz) oscillations can reflect information about working memory contents well past stimulus presentation^48^.

These within-condition decoding results validate that our signal of interest (information about the target orientation) was present in the data. For all further analyses, we use the voltage signal from the first 650 ms after stimulus onset to best capture the high temporal dynamics during perception and stimulus encoding, and we use the alpha-band signal from 650-2150 ms after stimulus onset to best capture the orientation signals during the delay period.

### Visual information is represented in between allocentric & retinocentric reference frames

Through head tilt, we can dissociate the allocentric and retinocentric reference frames. We hypothesized that visual representations (here: of orientations) may shift to a different reference frame when the head is tilted, compared to when the head is upright. A within-condition decoding approach cannot uncover such a shift, because any potential shift or bias is, by definition, going to be present in both the training and the testing set (Figure 2A). To overcome this limitation and uncover a potential shift in how orientations are represented, we used cross-generalized decoding: If target orientations are represented in a retinocentric reference frame, a decoder trained on head-upright trials would predict a 45º offset in decoded orientation when tested on head-tilted trials (after all, a vertical grating becomes diagonal on the retina after head tilt). Conversely, if target representations are allocentric, no such offset should be observed.

**Figure 2.**
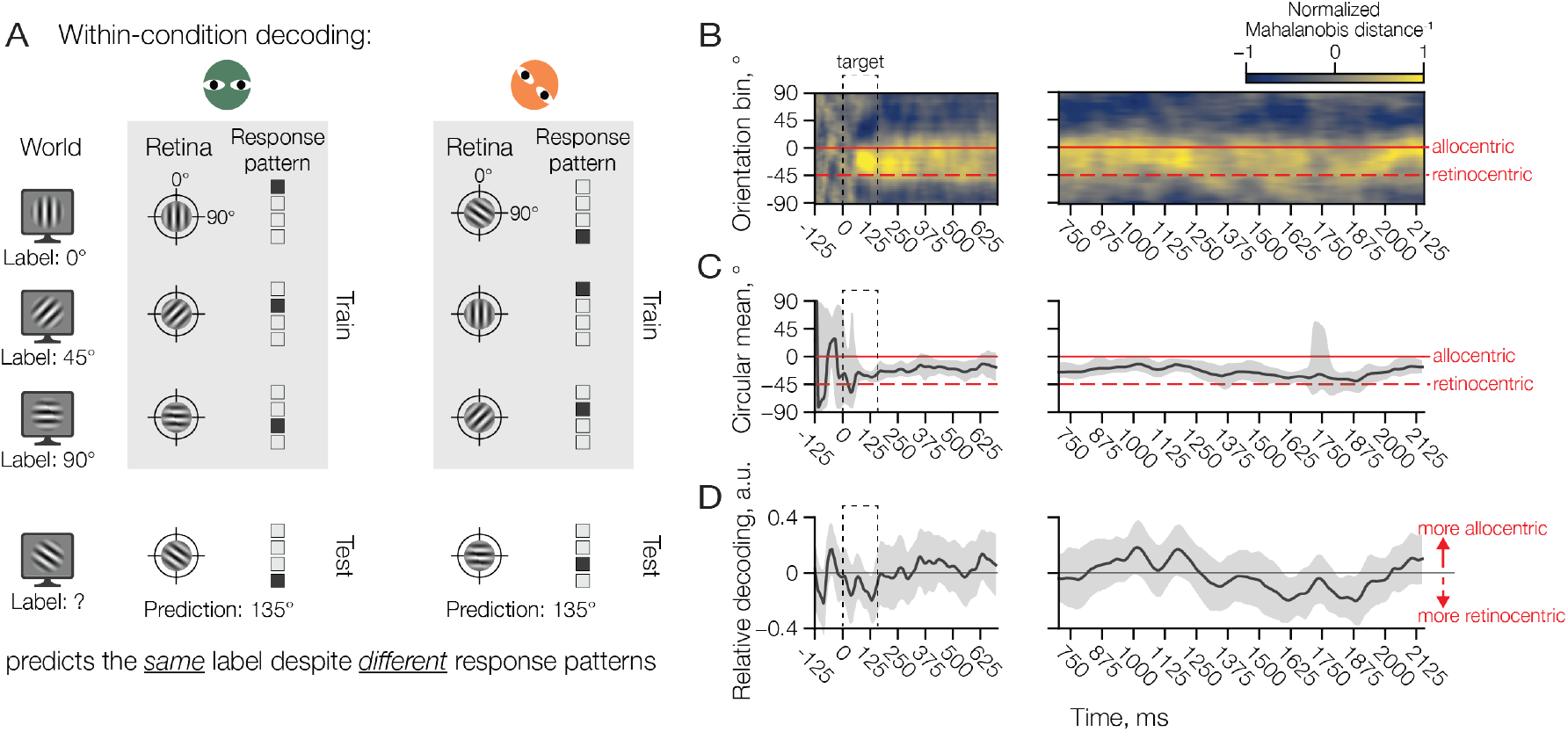
Cross-generalized decoding approach and results. **A**: A within-condition decoding approach cannot uncover a shift in reference frame when the head is tilted. To illustrate this point, we show a hypothetical shift in neural responses from trials with the head upright (leftmost panel) to trials with the head tilted (rightmost panel). When the head is upright, the allocentric (left column) and retinocentric (middle column) reference frames are aligned, and each orientation elicits a specific pattern of responses in the brain (right column). When training a decoder (top 3 rows), the label assigned to each orientation becomes associated with a specific pattern of responses, such that during testing a new label can be predicted from a new pattern of responses (bottom row). When the head is tilted, the allocentric orientation on the screen (left column) and the retinocentric orientation on the retinae (middle column) are no longer aligned. In this example, the pattern of brain responses (right column) has shifted relative to the patterns measured when the head was upright. However, since the decoder still associates the same assigned labels to this shifted pattern during training (top 3 rows), it will circularly interpret the pattern provided at test (bottom row) in the same way as before, thus predicting the same label as before. **B**: Cross-generalized decoding (trained on head-upright, tested on head-tilted, and vice versa), showing the average similarity (inverse Mahalanobis distance) of the target orientation to the binned orientations from the training set, plotted over time. Data are plotted such that the solid red line at 0º indicates the allocentric reference frame, while the dashed red line at -45º indicates the retinocentric reference frame. During the perceptual epoch (voltage signal, left) and the delay period (alpha signal, right) the target representation is most similar to orientations that are in between the allocentric and retinocentric reference frames. Note that, to avoid biasing our data towards participants with stronger cross-generalized decoding performances, participant-level Mahalanobis curves were z-scored before averaging across all participants. **C**: The circular mean of the population-level pattern similarity over time, with bootstrapped 95% confidence intervals. The circular mean can be interpreted as the peak similarity between the target orientation and orientations in the allocentric (dashed blue line) and retinocentric (dashed yellow line) reference frames over time. The circular mean remains between the two reference frames throughout the entire trail. **D**: Relative decoding (i.e., allocentric minus retinocentric) over time with bootstrapped 95% confidence intervals. At no point during the trial is relative decoding more allocentric or a more retinocentric, as indicated by the absence of clusters that significantly differed from 0 (based on a permutation test with cluster-defining threshold *p* < .05, corrected significance level *p* < .05).

Specifically, we trained a Mahalanobis-distance-based decoder on head-upright trials and tested it on head-tilt trials (and vice versa, see Methods). Smaller Mahalanobis distances indicate higher similarity between the pattern of responses evoked by a target orientation, and other possible orientations. Will this pattern similarity (i.e., inverse of the Mahalanobis distance) peak at an offset that indicates an allocentric or retinocentric reference frame? The Mahalanobis curves of the cross-generalized decoding, averaged across all participants, are shown in Figure 2B. Note how the target orientation is most similar to orientations somewhere in between the allocentric (solid red line) and retinocentric (dashed red line) reference frames, starting from early perception (left panel) and continuing throughout the entire delay (right panel).

We quantified this observed reference frame shift in two complementary ways. First, we calculated the circular mean of the population-level Mahalanobis curves, which resulted in direct estimates of the reference frame shift over time (Figure 2C). The target orientation was never significantly allocentric (0°) or retinocentric (-45º), but instead reflected a combination of the two reference frames. Already during perception the circular mean was -23.6º degrees (average voltage decoding over 0-650 ms; 95% bootstrapped CIs: -3.31º, -38.19º), and it remained in between the two reference frames throughout the trial, with a circular mean around -24.74º during the delay (average alpha-power decoding over 650-2150 ms; 95% bootstrapped CIs: -7.48º, - 38.59º). To verify that the reference frame did not shift rapidly from retinocentric to “in between” during the very earliest moments of perception, we used a highly robust decoder that also incorporates temporal dynamics of the voltage signals from the time period around the feed-forward sweep. Specifically, this decoder uses the concatenated data from 17 posterior channels and 5 timepoints from 50-100 ms after stimulus offset (in steps of 10 ms), to arrive at a single reference frame estimate for each participant. Even at this very early timepoint, a circular mean of -24.8º (95% bootstrapped CIs: -17.32º, -30.61º) indicated an orientation in between the two reference frames (Supplementary Figure 2A).

Second, we calculated a “relative decoding” metric by subtracting decoding performance in the retinocentric reference frame from decoding in the allocentric reference frame for each participant (Figure 2D). Thus, positive relative decoding indicates a more allocentric reference frame, negative relative decoding indicates a more retinocentric reference frame, and relative decoding around zero indicates equal evidence for both reference frames. In line with the results from the circular mean (Figure 2C), relative decoding was never significantly different from 0 (Figure 2D). Together, these results imply that when allocentric and retinocentric reference frames are dissociated via head-tilt, target orientations are represented not in any one reference frame, but in a combination of two reference frames together.

While there was little evidence to support orientation decoding from eye movements (outside one brief period from 875-1300 ms when the head was upright), we nevertheless applied the cross-generalization analysis also on signals from the EOG channels. If eye movements played a role in the neural results, we could expect peak decoding in between the allocentric and retinocentric reference frames here as well. Interestingly, insofar as orientation information was decodable from the eyes, it followed a predominantly allocentric reference frame (significantly so from ∼1400-1850 ms after stimulus onset; Supplementary Figure 1B-D). This implies that participants moved their eyes systematically relative to target orientations on the screen. While the combination of reference frames observed in the neural data could hypothetically be caused by a mixture of retinocentric brain signals and the allocentric eye movements, this seems unlikely – the allocentric contribution of eye movements was limited to a brief time window during the late delay, whereas the combination of reference frames observed in the neural signal was present throughout the entire trial.

### Representational geometry supports an “in-between” reference frame

We’ve shown that response patterns recorded with the head upright cross-generalize to response patterns recorded with the head tilted. This implies a shared, and partially shifted, coding scheme during head tilt. In addition to response patterns, we can look at the representational geometry during head tilt by using Representational Similarity Analysis (RSA). The representational geometry captures a lower dimensional format of a given set of stimuli, and is invariant to the underlying response patterns. This is because the geometry only captures the relationships between response patterns (e.g., how similar are responses to targets of 30º and 35º), such that also shifted patterns of responses could lead to the exact same geometry. Given that shared response patterns do not guarantee a shared geometry, does the representational geometry remain stable, or does shift with the head as well?

We calculated the representational similarities between responses to all target orientations during the first 50-100 ms after stimulus onset (see Methods for a detailed explanation) separately for the head-upright (Figure 3A) and head-tilted (Figure 3B) trials. In the representational similarity matrix (RSM) for head upright trials we observed a signature orientation anisotropy known from previous EEG studies^49^, namely, cardinal orientations were similar to themselves (i.e., 0º to 0º and 90º to 90º) but different from each other (i.e., 0º is different from 90º), while obliques were similar to themselves (i.e., 45º to 45º and 135º to 135º) but also similar to each other (i.e., 45º is similar to 135º)(Figure 3A). We used this signature orientation anisotropy to test what happens to the representational geometry during head tilt, which appears shifted (Figure 3B). To quantify this shift and its implications for to the reference frame during head tilt, we circularly shifted the head-tilted RSM five degrees at a time and calculated the mean absolute difference with the head-upright RSM at each step. Replicating our decoding results, this analysis indicates a geometry in between the allocentric and retinocentric coordinates when the head is tilted (Figure 3C).

**Figure 3.**
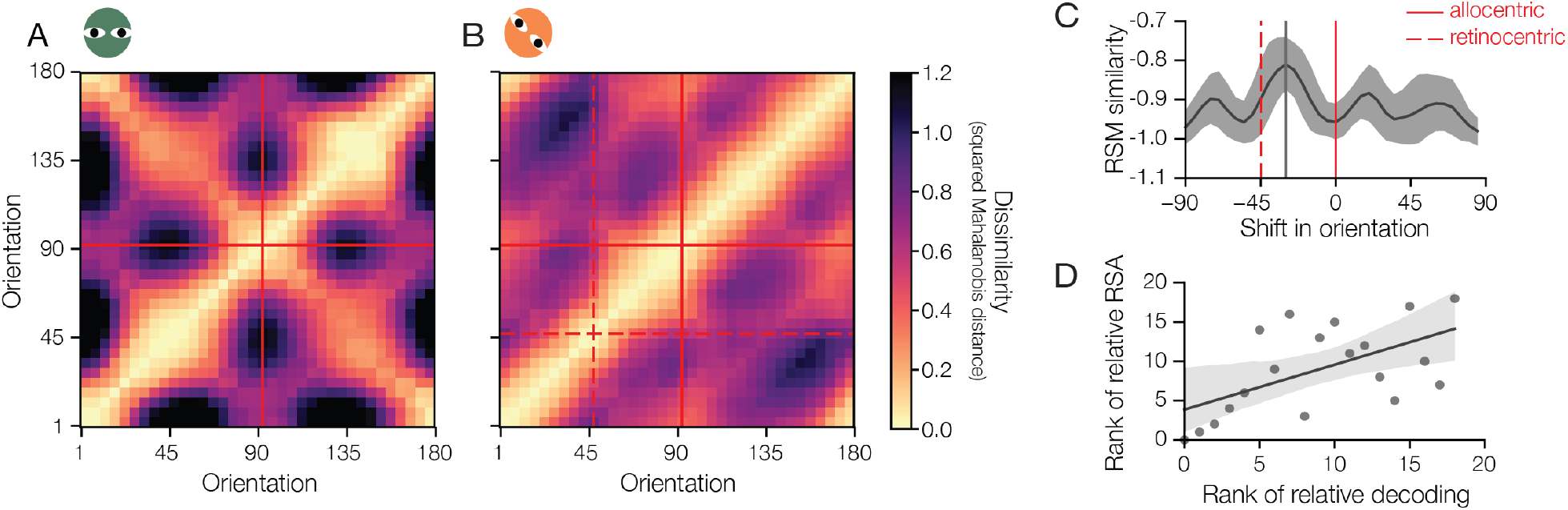
To assess whether head tilt also leads to a shift of the representational geometry, we ran a representational similarity analysis (RSA). The representational geometry of orientation across participants is plotted for head-upright trials (**A**) and head-tilted trials (**B**). Solid red lines denote the remembered orientation in the allocentric reference frame, dashed red lines denote the remembered orientation in the retinocentric reference frame. Note that in head-upright trials, allocentric and retinocentric reference frames are equivalent, so only the allocentric line is visible. **C:** To more precisely quantify the change in the head-tilted trials, we continuously shifted the head-tilted RSM in steps of 5 degrees and calculated the similarity between each shifted head-tilted RSM and the original head-upright RSM. We calculated the mean absolute difference between the cells of the two RSMs for each shift for each participant. The group-level means are plotted with bootstrapped 95% confidence intervals. The highest similarity occurs when the head-tilted RSM is shifted in between the allocentric (solid red line) and retinocentric (dashed red line) reference frame. **D:** To test whether the individual differences we found in the cross-generalized decoding align with the individual shifts quantified in **C**, we correlated the estimated amount of shift in head-tilted trials based on the RSA and the cross-generalized decoding. The ranked amounts of shift are plotted together with a linear regression model fit with a bootstrapped 95% confidence interval over the beta weight (equivalent to correlation coefficient).

Finally, we wanted to directly compare our decoding and RSA results. To this end, we calculated a “relative RSA” for each participant and correlated this with relative decoding. The “relative RSA” was calculated by centering the head-tilted RSM in allocentric and retinocentric coordinates, and taking the mean absolute difference between the head-upright RSM. The difference between the two resulting values reflects the degree to which a participant’s geometry was more allocentric or retinocentric. We observed a positive correlation between this “relative RSA” and relative decoding (Figure 4D, Spearman correlation: *r*_(17)_ = .572, *p* = .0105).

**Figure 4.**
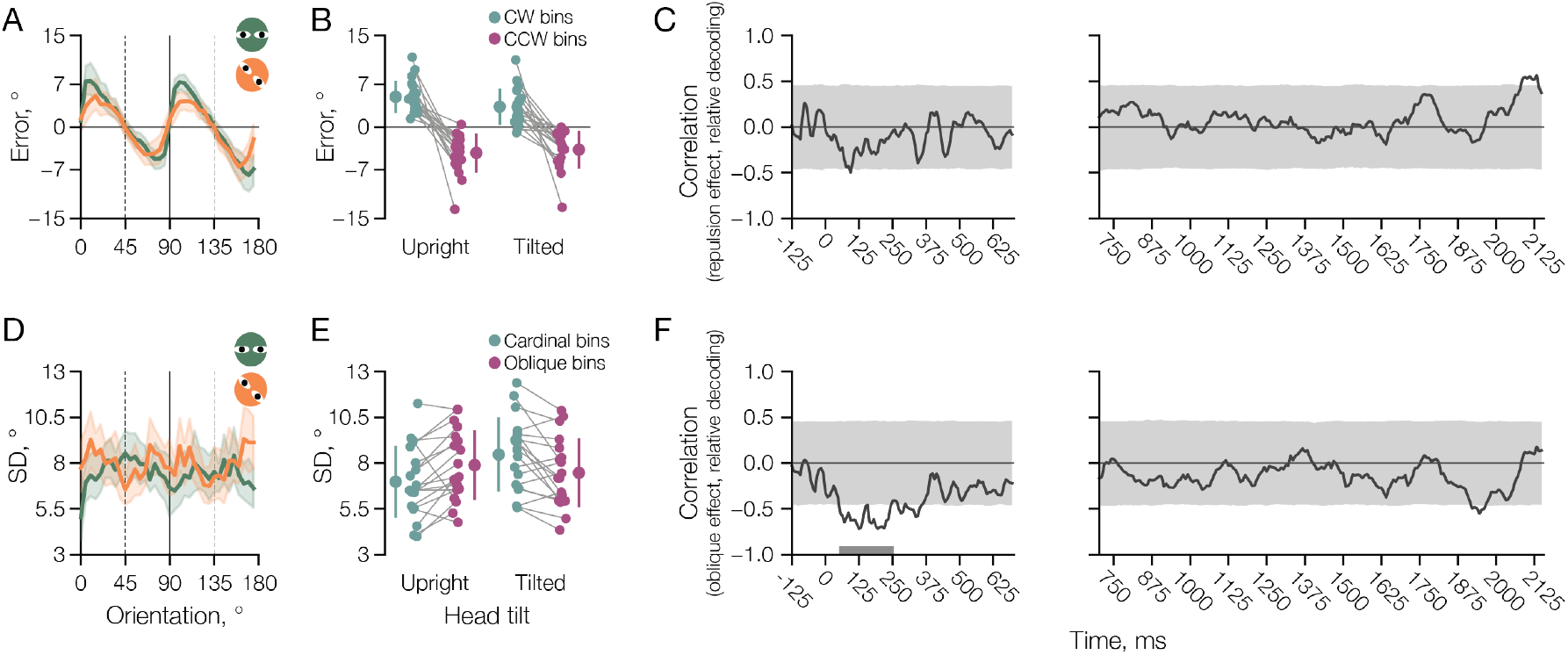
Repulsion and oblique effects, and their correlations with relative cross-generalized decoding. **A-B:** Responses to target orientations near allocentric cardinal axes were biased away from those axes in both head-upright and head-tilted trials: For targets that were clockwise of a cardinal orientation, recall errors were biased in a clockwise direction (error > 0), and vice versa for counterclockwise orientations (error < 0). Thus, the repulsion effect is anchored to an allocentric reference frame. In **A** the signed recall error is plotted as a function of the target orientation (in allocentric coordinates) across all participants. Shaded areas indicate bootstrapped 95% confidence intervals. In **B** this repulsion is quantified for individual participants by dividing the orientation space into two bins – one clockwise (teal) and one counterclockwise (purple) of allocentric cardinal orientations. The repulsion effect was somewhat more pronounced when the head was upright (F(1,18) = 9.09; p = 0.0074), but present irrespective of head tilt (F(1,18) = 48.56; p < 0.001). **C:** Correlation between the strength of the repulsion effect and relative decoding, plotted over time. No clusters were significantly correlated. Shaded areas represent the permuted null-distribution of the correlation coefficients. **D-E:** The oblique effect shifted along with the tilt of participants’ heads: The circular standard deviation (SD) was lower around allocentric cardinal orientations compared to allocentric obliques when the head was upright, and was higher around allocentric cardinal orientations compared to allocentric obliques when the head was tilted (meaning these obliques become cardinals in retinocentric coordinates). Thus, the oblique effect is anchored to a retinocentric reference frame. In **D** the circular SD is plotted as a function of the target orientation (in allocentric coordinates) across all participants. In **E** the oblique effect is quantified for individual participants by dividing the orientation space into allocentric cardinal (teal) and allocentric oblique (purple) bins. Responses for allocentric cardinal orientations were more precise (smaller SD) with the head upright, and less precise (larger SD) with the head tilted (F(1,18) = 54.56; p < 0.001), and responses were somewhat more precise with the head upright (F(1,18) = 34.76; p < 0.001). **F:** Correlation between the strength of the oblique effect (allocentric cardinal-minus-oblique) and relative decoding, plotted over time. The strength of the oblique effect significantly correlated with relative from approximately 50-250 ms after target onset (so during perception). This implies that that participants with a stronger oblique effect in head tilted trials (i.e., relatively low precision for retinocentric cardinals compared to retinocentric obliques) encode orientations in a more retinocentric reference frame. The teal bar just above the x-axis denotes a significant decoding cluster (permutation test, n = 19, cluster-defining threshold *p* < .05, corrected significance level *p* < .05).

Taken together, the results from our decoding and RSA analyses show that when a person’s head is tilted, both the neural coding scheme (i.e., response patterns) and the coding geometry indicate that visual orientations are represented in a combination of allocentric and retinocentric reference frames during early perception, and that individual differences in the amount of shift persist across these two different measures of neural coding.

### Individual differences, and the correlation between decoded reference frame and strength of the oblique effect

Thus far we’ve demonstrated that, on average, visual information is represented “in between” an allocentric and retinocentric reference frame when people tilt their heads. However, one thing we have not yet discussed are inter-individual differences. To obtain a robust estimate of each individual participant’s decoded reference frame, we first tested for inter-individual differences at the neural level by using results from the same robust spatiotemporal decoder applied to the very earliest moments of perception (Supplementary Figure 2). For about half of participants (n = 10), the difference between allocentric and retinocentric accuracy was not significantly different from zero, in line with an “in between” reference frame. However, for the other half of participants the reference frame used was primarily allocentric (n = 4) or retinocentric (n = 4) in nature (Supplementary Figure 3). This establishes clear inter-individual differences in the reference frame decoded from our EEG data.

To explore individual differences in behavior, we turned to two known effects in orientation response errors - the oblique effect and the repulsion effect. First, we replicated previous behavioral findings^35^, showing that the repulsion effect is tied to the allocentric reference frame (i.e., repulsion relative to world-centered vertical and horizontal, Figure 5A), while the oblique effect is tied to the retinocentric reference frame (higher resolution for retina-centered cardinals, Figure 5D). Together with these group effects, we saw clear individual differences in the strength of the repulsion effect (Figure 5B) and oblique effect (Figure 5E). The strength of the repulsion and oblique effects for each individual participant were defined as the difference between cardinal and oblique bins (cardinal-minus-oblique) on trials where the head was tilted.

Having established individual differences at both the neural and behavioral level, we ran an exploratory analysis to investigate their relationship. We correlated the strength of the repulsion and oblique effects for each individual participant with their decoded evidence for allocentric or retinocentric reference frames (i.e., relative decoding) over time. While the repulsion effect did not significantly correlate with the decoded reference frame (Figure 5C, no significant clusters found), the strength of the oblique effect was negatively correlated with the decoded reference frame for a significant cluster of time during presentation of the target orientation, from approximately 50 to 250 ms (Figure 5F). Note that a *larger* relative decoding value indicates a more allocentric reference frame. Also, a *larger* cardinal-minus-oblique difference indicates a stronger oblique effect. This is because the oblique effect tilts along with the head, such that precision is relatively higher at the allocentric oblique (which becomes a retinocentric cardinal with head tilt) compared to the allocentric cardinal (a retinocentric oblique with head tilt). Thus, this negative correlation suggests that participants who show a stronger oblique effect (i.e., have relatively low precision for retinocentric cardinals compared to retinocentric obliques) encode orientations in a more retinocentric reference frame. We suggest that using real-world allocentric cues (like the edges of the computer screen) probably helps increase behavioral precision when reporting a vertical or horizontal orientation (i.e., allocentric cardinals). However, when someone relies on a more retinocentric reference frame they probably use these allocentric cues to a lesser extent, which might result in relatively imprecise reports for vertical and horizontal orientations. This could explain why a more retinocentric reference frame was associated with relatively lower behavioral precision for allocentric cardinals.

## Discussion

The human brain could represent visual information in a reference frame anchored to the eyes (retinocentric) or the outside world (allocentric). In this study, we used head tilt as an experimental manipulation to dissociate these two reference frames, and consistently found that visual information for orientation was represented via a combination of retinocentric and allocentric reference frames, both during perception and memory. Using EEG recordings in humans, this finding aligns with work in non-human primates which has also shown “in between” coding at the level of single neurons.

Already from the earliest decodable timepoints, when the target orientation was still present on the screen, cross-generalization between head-upright and head-tilted trials showed an orientation representation in between retinocentric and allocentric coordinates (based on the analysis of voltage signals). This persisted once the target orientation was no longer on the screen and was held in mind until the end of the delay (based on the analysis of alpha signals^48^). Such a combined representation during working memory delay suggests that, even when the retinal input stops, the reference frame in which visual information is represented persists. This overlap between perception and during memory could be due to the recruitment of visual cortex to represent working memory content^39,51,52^. At no point during the trial was there significant evidence for one or the other reference frame being more prevalent. Importantly, significant within-condition decoding of orientation information throughout the trail strongly suggests that this finding is not due to lack of signal in either reference frame. Moreover, the ability to cross-generalize between head upright and head titled trials demonstrates that the brain uses a shared coding scheme to represent orientations in both cases, and that this coding scheme partially shifts with head tilt. In addition to the coding scheme (i.e., measured via response patterns), also the representational geometry (i.e., measured via RSA) is shared and shifted with head tilt.

To our knowledge, this is the first direct evidence that visual representations in the human brain can deviate from a purely retinocentric format already at the earliest stages of perception in humans. Previous studies of human neural representations have consistently demonstrated that, during perception, visual representations are represented in a retinocentric (i.e., retinotopic) reference frame^1,3^. This difference between previous research and the work presented here is likely due to the manipulation used to dissociate retinocentric and allocentric reference frames: Previous studies have used gaze manipulations, which may not be representative for uncovering reference frame transformations^12^, while we used head tilt. This highlights a potentially important difference between eye movements and head or body tilts: to account for eye movements, one needs to rely on efferent motor commands from eye muscles, which are transient and may themselves operate in a retinotopic reference frame. In contrast, to account for head tilt, one can rely on gravity signals processed by the vestibular system, which (on earth) are constant, omnipresent, and unlikely to be maintained in a retinocentric format.

As already mentioned, the “in between” representations we see in our study align with previous research in non-human primates that have used head tilt to investigate egocentric and allocentric frame integration and showed evidence for a combined representation. Most notably, Rosenberg and Angelaki^27^ demonstrated that head-and-body tilt causes the neurons in the monkey central intraparietal cortex (CIP) that respond to a specific planar tilt to shift towards a combination of allocentric and retinocentric signals. But also in the primary visual cortex (V1), neuronal responses can be differentially affected by heading directions^50^. While changes in heading direction are not identical to changes in head or body tilt, they are often aligned because the head and body are typically oriented in the direction of an animal’s movement.

Of course, with EEG we cannot exactly pinpoint the origins of the signals measured, and cannot draw any firm conclusions about the involvement of specific regions, such as the intraparietal or primary visual cortex. Characteristics of neurons may vary significantly between these different regions of the visual hierarchy. Thus, the intermediate representation found in this study could be a combination of retinocentric representations in e.g., the primary visual cortex, and allocentric representations in e.g., in the parietal cortex. The influence of head tilt on neural representations is challenging to study in humans due to methodological limitations such as the lack of spatial resolution (e.g., in EEG or fNIRS). Methods with superior spatial resolution (e.g., fMRI or MEG) generally do not allow for head movements. Nevertheless, our results align well with the beforementioned electrophysiology work, and are consistent with a combined reference frame for visual processing in the visual cortex more broadly construed.

In addition to not being able to pinpoint the involvement of various visual areas, the non-invasive EEG signal is also noisy and requires averaging for robust interpretation. This means we cannot distinguish between the possibility that there is one consistently maintained “in between” reference frame on every single trial, or the possibility that multiple heterogeneous reference frames, ranging from retinocentric to allocentric, are engaged across different trials but result in a single “in between” average. Electrophysiological data from monkeys points towards the latter: when a monkey’s head and body are tilted, tuning functions of parietal neurons shift heterogeneously, resulting in a continuum of representations from retinocentric, to intermediate, to allocentric reference frames^27^.

Adequately representing visual input is a challenging task – the input itself is necessarily retinocentric, and multiple sources of eye, head, or body movements may affect the relationship between this retinocentric signal and the world that is its source. Our study shows that, during head tilt, these factors are not combined in an “all-or-nothing” way, with potentially useful information being discarded. Instead, both retinocentric and allocentric information are already encoded during perception, and information about these two reference frames is maintained during memory, after the retinal input has stopped.

## Methods

### Participants

A total of 25 volunteers (13 female) between the ages of 18 and 32 years (mean = 22.12, SD = 4.03) were recruited to participate in this study. Note that due to a technical error the ages of S07 and S08 were not recorded (and not included in the mean and SD age). Of these, 19 participants were included in the final analyses due to several participants dropping out (N = 3), technical issues during data recording (N = 2), and a serious neurological condition (N = 1). The study was conducted at the University of California San Diego, and approved by the local Institutional Review Board. All participants provided written informed consent, had normal or corrected-to-normal vision, and received a monetary reimbursement of $15 per hour.

### Stimuli & procedures

Stimuli were generated using MATLAB 2016a and the Psychophysics toolbox^54,55^ on a computer running Ubuntu 16.04 LTS. Stimuli were viewed on a luminance-calibrated CRT monitor (1600 × 1200 resolution, 100 Hz refresh rate) against a mean grey background (128 cd/m^2^) from a distance of 50 cm in an otherwise darkened room. Stable head and eye positions were aided by a custom chinrest, and a centrally presented white fixation circle with inner- and outer-radii of 0.125º and 0.25º, respectively. Participants were instructed to fixate throughout. The stimuli used as memory targets were sinusoidally modulated gratings with a spatial frequency of 0.65 cycles/º, 20% Michelson contrast, and randomly jittered phase (1-360º). These gratings were configured in a donut-shaped circular aperture with a 0.75º and 12º inner and outer radius, respectively, and smoothed edges (0.245º Gaussian kernel; SD = 0.136º). Grating orientations were uniformly sampled from 5º to 180º in 5º steps, meaning that, on a given trial, the target orientation could be 5º, 10º, 15º, and so on, up to 180º. These 36 possible target orientations were presented in randomly shuffled order across trials. Note that, throughout the paper, we report orientations in the allocentric reference frame – the distal stimulus orientation as it was presented on the display. The recall dial consisted of two short (0.25º) white lines that rotated around the fixation dot as if they were part of one single line (0.25º wide) of which the central-most 12º were not shown.

On each trial of the memory task, participants were presented with a target grating for 150 ms, and they remembered its orientation over a 2000 ms delay. After the delay, participants saw a randomly oriented line (the recall dial) that they could rotate to match the orientation in memory by moving the computer mouse. Once participants were satisfied with their response, they clicked a mouse button to continue to the next trial in an unspeeded manner. Inter-trial intervals were 500 ms + 50 ms of random temporal jitter. Critically, on half of the trials, participants’ heads were tilted by 45º relative to upright while performing this task. Specifically, on every run the tilt of a participant’s head was alternated in 4 blocks of 72 trials per block. In other words, a participant would start a run with their head in a given position (e.g., upright) and would complete 72 trials, after which they received the instruction to change their head tilt (e.g., to 45º clockwise) for the next 72 trials, and so forth, until 4 blocks were completed. Participants did not tilt their heads by themselves. Rather, the experimenter would enter the EEG chamber in between blocks to rotate the custom-made chinrest (so designed that the center of rotation would be the nasion of the participant), ensuring the appropriate head position for the next block. Per EEG recording session, participants completed 3 runs of this memory task (of 288 trials per run), which meant that they completed a total of 432 trials per head-tilt condition per recording session. The direction of head tilt was counterbalanced across subjects, with 9 participants tilting clockwise, and 10 participants tilting counterclockwise.

In total, each participant took part in at least three EEG recording sessions. The central electrode (Cz) was placed exactly in the middle of nasion to inion in every session. If one of the sessions finished early for technical or logistical reasons, the participant came in for a fourth recording session. Given that a typical session consisted of 3 runs of the memory task, we collected a total of 2592 trials (1296 per head-tilt condition) across all sessions. While this was the case for 13 out of 19 participants, for 3 participants one extra run of the memory task was available (a total of 2880 trials), and for 1 participant one run fewer was available (a total of 2304 trials). For two other participants, EEG recordings from one session were rendered unusable by a CMS/DLR issue in the recording system, and only the two intact sessions of data were analyzed (a total of 1728 total trials). In addition to the memory task described above, all participants also performed a perceptual task in which they viewed the same grating stimuli as in the memory task, but had to detect a brief contrast change (without a memory delay or recall phase). Data from this perceptual task were not used in the analyses of the current study beyond the ICA preprocessing stage.

### Electroencephalography (EEG) acquisition and preprocessing

Electroencephalography (EEG) data were recorded using a 64+8 channel BioSemi ActiveTwo system (Amsterdam, The Netherlands) at a sampling rate of 1024 Hz. The 64 scalp electrodes were laid out according to the extended international 10-20 system. Of the 8 external electrodes, 2 reference electrodes were placed at the mastoids, and 6 external electrodes were used to monitor eye movements and blinks via Electrooculography (EOG). To this end, three electrodes were placed around each eye: below, above, and near the outer canthi of the eye. All EEG data were referenced online to the BioSemi CMS-DRL reference, and we aimed to keep all offsets from the reference below 10 μV.

EEG data were preprocessed and analyzed using selected functions from MNE-Python^56^, SciPy^57^, pingouin^58^, the FieldTrip extension in Matlab^59^, and custom Python and Matlab scripts. Before preprocessing, we first ran an independent component analysis (ICA). For this ICA, the data were concatenated across all runs of each EEG recording session and bandpass filtered (1 Hz high-pass and 40 Hz low-pass Hamming windowed sinc FIR filter). Periods during which the participants changed their head tilt between blocks were marked as breaks using the MNE-Python annotate_breaks() function with additional visual inspection and manual adjustments where needed. Noisy and malfunctioning electrodes were flagged for exclusion at this point too, based on visual inspection of the filtered concatenated runs. An average of 5.15 electrodes were flagged for exclusion per recording session (range 0-13 electrodes) and excluded from all further analyses. The data were down-sampled to 256 Hz, and then decomposed with FastICA using the preprocessing.ICA() function from MNE-Python^60^. The resulting independent components were inspected visually. Canonical eye movement components (blinks and saccades) were marked for exclusion. One subject’s ICA results contained a strong heartbeat component that was also marked for exclusion. These various outcomes of the ICA were used at several stages of data preprocessing, as described next.

To prepare the EEG data for the main analyses, raw data were preprocessed for each experimental run independently. The data were bandpass filtered (0.1 Hz high-pass and 40 Hz low-pass Hamming windowed sinc FIR filters), and flagged electrodes and ICA components from the corresponding session were removed from the data. The data were then referenced to the average of all remaining scalp electrodes, and the flagged electrodes were interpolated using a spherical spline method as realized in the MNE interpolate_bads() function. The data were epoched –200 ms to 2150 ms relative to the target grating onset. Next, we sub-selected 17 posterior electrodes (P7, P5, P3, P1, Pz, P2, P4, P6, P8, PO7, PO3, POz, PO4, PO8, O1, Oz, and O2), the same electrodes as in previous studies of visual working memory using EEG (e.g. Wolff et al 2015, 2017), for all subsequent preprocessing steps and analyses. We visually inspected the resulting epochs from these 17 channels using the FieldTrip ft_rejectvisual function, which shows summary statistics across channels, epochs, or both. Based on this visual inspection, we removed outlier epochs from all further analyses. In the head-tilt runs, an average 3.36% (+3.87%) of epochs were rejected per participant, with a maximum of 18.33% for one participant. In the head-upright runs, an average 3.08% (+2.67%) of epochs were rejected per participant, with a maximum of 11.81%.

The goal of our preprocessing pipeline was to obtain two types of data for each timepoint of the EEG signal for further time-course decoding analyses – the broadband voltage signal and the alpha-band power signal. First, for the time-course decoding analyses on the broadband voltage signal, each epoch was baselined using the average signal from -200 ms to 0 ms before stimulus onset. After baselining, each electrode’s data were demeaned, independently at each timepoint, by subtracting the average voltage for all electrodes included in the analyses at the corresponding timepoint. Subsequently, the time-courses of each epoch and channel were smoothed using the SciPy gaussian_filter1d() function with a Gaussian kernel of SD = 16 ms. Finally, the data were downsampled to 128 Hz. Second, to isolate the signal related to alpha-band oscillations, we computed the spectral power of the EEG signal from 8 to 12 Hz in steps of 1 Hz using Hanning tapers with sliding time-windows (in steps of ∼7.8 ms, or 128Hz) of 3 cycles per frequency using the FieldTrip ft_freqanalysis() function. The obtained alpha power estimates for all frequency bands were subsequently averaged, resulting in a single alpha power estimate at each timepoint. For the time-course decoding analyses, alpha power was further demeaned across electrodes in the same manner as for the broadband voltage. Finally, for the control analysis using signals obtained from the eye channels, the six unipolar EOGs were reconfigured into one bipolar HEOG channel (using the 2 unipolar electrodes at the outer canthi of the eyes) to track horizontal eye movements, and the average of two bipolar VEOG channels (using the 2 sets of unipolar electrodes above and below each eye) to track vertical eye movements.

### Analyses

#### Orientation decoding

To decode orientation, we computed the similarity in EEG response patterns to different orientations^45,61^ at different time points. To this end, we used the inverse of Mahalanobis distance as a measure of pattern similarity, which captures the assumed increase in similarity between more similar orientations.

First, all trials were assigned to the closest of 12 orientation bins centered at 0°, 15°, 30°, and so on up to 165°. We then split the data into training and testing sets using a stratified k-fold cross-validation approach with k = 12 folds. Within each fold, 11/12^th^ of the trials were used as the training set, and the remaining 1/12^th^ was used as the testing set. Across all 12 folds within one repetition, every 1/12^th^ of the data served as the testing set once. In case of non-uniformities in the distribution of trials across orientation bins in the training set (due to, e.g., artifact rejection), we randomly subsampled trials in the training set so that each orientation bin had an equal number of trials, to avoid bias in the decoding. On each fold, the subsampled training trials were used to estimate the noise covariance matrix using a shrinkage estimator^62^. To compute the covariance matrix, we first estimated the residual of the orientation signal by removing the average signal of each orientation bin from the individual trials in that bin. Note that this was only done for the estimation of the noise covariance. Also on each fold, the subsampled training trials of each orientation bin were averaged and convolved with a half-cosine basis set raised to the 11^th^ power to make use of the continuity of the orientation space and pool information across similar orientations^63^. These smoothed bin means, together with the noise covariance matrix, were used to calculate Mahalanobis distances between the orientation bins of the training data and the trials of the testing data.

Specifically, for a single testing trial and a single training bin, the Mahalanobis distance, *d*_*M*_, between the testing trial 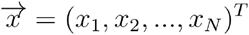, where *N* = 17 is the number of posterior electrodes used for analysis, and the multivariate probability distribution that is the training bin *Q* which is parameterized with mean 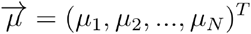 and the pseudo-inverse of the covariance matrix *S*^−1^ was calculated as:

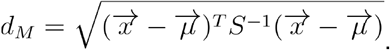

Calculating the Mahalanobis distance between a single test trial and each training bin resulted in 12 Mahalanobis distance values per test trial in the testing set that were normalized by mean-centering. This entire cross validation procedure was repeated 20 times, with a new stratified 12-fold cross-validation split randomly generated on each repetition. Thus, within the one binning scheme already described thus far, the 12 Mahalanobis distance values per time point and trial were estimated 20 times. These values were averaged across the 20 repetitions.

We then repeated the same process for two alternative orientation binning schemes: one with bins centered at 5°, 20°, 35°, and so on up to 170°, and one with bins centered at 10°, 25°, 40°, and so on up to 175°. Across the three total orientation binning schemes, 36 Mahalanobis distances were acquired (from equidistant orientation bins, spaced 5º apart). Each of these distances represents the dissimilarity between the multivariate EEG patterns associated with each of the 36 total orientation bins, and the multivariate EEG pattern associated with one of the 36 possible target orientations on a given time point and trial. In case of an orientation-specific signal in the data, the Mahalanobis distance will be smaller for an orientation bin close to the target orientation, and larger for orientation bins further away from the target orientation. To quantify this orientation-specific signal, the 36 Mahalanobis distances for each time point were shifted such that the orientation differences between the test trial and training bins ranged from –90º to +85º, with the shared orientation (i.e., 0º difference) indicating the center. Since distance is a measure of pattern *dissimilarity*, we took the inverse of the distance as a measure of similarity. This means that for each time point that contains orientation-specific information, a “Mahalanobis curve” with an inverted-U shape is anticipated (i.e., the highest similarity in the center, where the orientation bin matches the target orientation). Conversely, Mahalanobis distances are not expected to systematically differ between orientation bins on time points without orientation information (i.e., a flat Mahalanobis curve). To quantify the amount of orientation information at different time points, we calculated the cosine vector mean of the Mahalanobis curve at each time point for each participant. This metric is used as our measure of orientation decoding accuracy.

The decoding approach described above was applied within each condition (head upright and head tilted) for both broadband voltage and alpha power signals. However, to test whether neural representation of orientation is more allocentric or retinocentric, and to quantify a possible shift over time, it is necessary to do cross-generalized (i.e., across condition) decoding. A decoder, ultimately, just receives response patterns and their corresponding labels. In our case, these response patterns are broadband voltage or alpha-band power across multiple EEG channels, and the labels are orientation values in the allocentric reference frame. During training, the decoder estimates some relationship between response patterns and labels. During testing, the decoder is presented with new response patterns to assign labels to. This label assignment is restricted to the relationship between labels and response patterns learned during training, which does not allow us to uncover a representational shift. To illustrate, imagine that at some time point in the head tilt condition the representation of a given orientation (say 0º in allocentric coordinates) has shifted to somewhere in between allocentric and retinocentric (so in between 0º and 45º, e.g., 22.5º). Yet, if we train and test within this time point and within head tilted trials, the only answer the decoder can give us is that the orientation was 0º. After all, the decoder has learned that the observed pattern of responses (shifted or not) is associated with the label “0º”. Cross-generalized decoding can circumvent this problem, by training on response patterns in one reference frame (e.g., when the head is upright), and seeing how these patterns cross-generalize to another reference frame (e.g., when the head is tilted). In case of a purely allocentric coding scheme, the pattern of responses to a vertical orientation viewed with the head upright will not change when someone tilts their head (even when that orientation is now 45º on the retinae). However, in case of a purely retinocentric coding scheme, the pattern of responses to a vertical orientation on the screen, viewed with the head tilted (i.e., 45º on the retinae), should look more like that of a 45º orientation on the screen viewed with the upright (i.e., also 45º on the retinae).

For cross-generalized decoding, we used trials on which the head was upright as the training set, and trials on which the head was tilted as the testing set, and vice versa. Cross-generalized decoding was performed analogously to what was described for the cross-validation approach, with the exception that no folding was needed (and was therefore skipped). Instead, results were averaged across the two cross-generalized decoders (one trained on head-upright trials and tested on head-tilted trials, and one trained on the head-tilted trials and tested on head-upright trials). We still randomly subsampled trials in the training set so that each orientation bin had an equal number of trials. This random subsampling (and hence the decoding) was repeated 20 times per binning scheme and per decoder, and the resulting Mahalanobis distances were averaged across these 40 repetitions (20 per decoder), same as in the cross-validated analysis. The 36 Mahalanobis distances for each time point and each trial were shifted such that they ranged from –90 degrees to +85, with a difference of 0 representing the same allocentric orientations, and a difference of -45 representing the same retinocentric orientations. This also means that, in order to align the decoding results across head tilt directions, we flipped the Mahalanobis curves for the participants who tilted their head counterclockwise. To avoid biasing the estimated shift towards participants with generally stronger decoding, the trial-averaged Mahalanobis curves of each participant were z-scored before calculating the population-level curves profiles.

To assess the results of the cross-generalized decoding, we used two complementary metrics. Firstly, we calculated the circular mean of the cross-generalized Mahalanobis curve averaged across subjects (i.e., at the population level). This metric is referred to as “population circular mean” throughout. Secondly, we calculated participant-level differences between the allocentric and retinocentric cosine vector means of the Mahalanobis curve. Specifically, we calculated the allocentric decoding accuracy (cosine vector mean between the Mahalanobis curve and a cosine centered on 0°), and the retinocentric decoding accuracy (cosine vector mean between the Mahalanobis curve and a cosine centered on -45°). We then subtracted retinocentric decoding accuracy from the allocentric decoding accuracy for each participant, such that a higher value indicates a more allocentric reference frame, a lower value indicates a more retinocentric reference frame, and a value of zero indicates equal evidence for both reference frames. This metric is referred to as “relative decoding” throughout. To examine the possible role of eye movements, the same analysis was repeated on the two EOG bipolar channels (horizontal and vertical).

#### Early perception decoding

Our main decoding analyses as described above, were performed at each timepoint separately, only including different electrodes as features – as is typical for this kind of analysis. However, temporal dynamics of the evoked responses in EEG can be informative as well, and pooling over these temporal dynamics along with dynamics across electrodes can improve decoding accuracy (at the cost of temporal resolution)^45,64,65^. In addition to being more sensitive, this analysis provides the most stringent test of what reference frame might be used during the initial feedforward sweep of visual processing – during the very earliest perceptual time window, from 50 ms to 100 ms. This will help establish if visual representations are retinocentric during the feedforward sweep, or if a shift in the decoded reference frame is present already during early perception. This cross-generalized analysis differs from what was previously described in the following ways: For each trial and channel separately, we subtracted the mean activity level within that time window to isolate the stimulus-evoked neural signal. In line with previous studies^45^, the data were then downsampled from original 1024 Hz to 100.24 Hz by taking an average every 10 ms to decrease computational demands. This resulted in 5 time points for each of the 17 posterior channels, which each time point of each channel used as a separate feature in the analysis (i.e., 85 features in total). The same spatiotemporal data were used for representational similarity analysis (RSA, see below).

To test whether the decoded reference frame was consistent within individual participants, we used these early perception decoding results, and again calculated relative decoding (allocentric minus retinocentric). We used a permutation test for each participant and each decoder (i.e., trained on head upright and tested on head tilted, and vice versa), to test whether the relative decoding accuracy was significantly different from 0. If a participant’s relative decoding was significantly above 0 on both decoders, they were considered to represent orientation in an allocentric reference frame. If a participant’s relative decoding was significantly below 0 on both decoders, they were considered to represent orientation in a retinocentric reference frame. Otherwise, they were considered to use a combination of reference frames.

#### Representational Similarity Analysis (RSA)

Representational Similarity Analysis (RSA) allows us to look at the representational geometry of orientation stimuli – how similar the representation of each orientation is to every other orientation. This geometry can be non-uniform^49^. Here, we rely on this non-uniformity to investigate a potential shift in representational geometry from head-upright to head-tilted trials during early perception (50-100 ms). To this end, we use a cross-validated RSA with inverse squared Mahalanobis distance as the similarity measure^66^. We used the same stratified 12-fold cross-validation approach with 3 binning schemes as described under “Orientation decoding”. Within each cross-validation fold, we subsampled the data in both the training and the testing set and calculated the squared Mahalanobis distance for the difference in each pair of orientation bins between the training and testing set. Specifically, for a single pair of orientation bins A and B, with trial-averaged multivariate response patterns 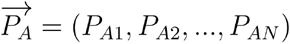 and 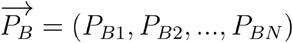, the squared Mahalanobis distance,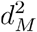, was calculated as: 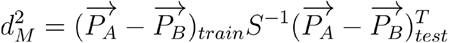, where *S*^−1^ is the pseudoinverse of the covariance matrix estimated from the training data, and `N` is the number of features used for analysis.

For each participant, the RSA resulted in two representational similarity matrices (RSMs), one for the head-upright trials, and one for the head-tilted trials. To avoid biasing the group-level effects towards participants with generally stronger decoding, we z-scored participant level RSMs.

To visualize the shift in geometry when the head was tilted relative to when the head was upright, we continuously shifted the head-tilted RSM in steps of 5° and calculated the mean absolute difference between head-tilted and head-upright RSMs at each step. Separately, to quantify the shift for each participant in the head-tilted RSM compared to the head-upright RSM, we centered the head-tilted RSM in both allocentric and retinocentric coordinates and calculated the mean absolute differences with the head-upright RSM (i.e., a “relative shift”). The difference of these two values reflects whether, for each participant, the representational geometry of the head-tilted RSMs is in more allocentric or more retinocentric coordinates, compared to the head-upright RSMs. Finally, to compare the results of the RSA to those from the cross-generalized decoding, these RSM difference values were correlated (Spearman’s rank correlation) with the relative decoding (allocentric minus retinocentric) obtained from the same early perception period (50-100 ms).

#### Behavioral effects, and their relationship to neural effects

Previous research has suggested that two canonical types of orientation recall errors, the oblique effect and the repulsion effect, behave differently during head tilt^35^. To test whether we could reproduce this in our data, we calculated the circular mean and standard deviation of the recall errors using the circmean and circstd functions from SciPy. At the group level, we calculate the circular mean and standard deviation at every possible target orientation (from 5º to 180º in steps of 5º) across all participants, while at the individual subject level we calculate the circular mean and standard deviation on binned data as described below.

To quantify the oblique effect at the level of individual participants, we divided the orientation space into two equally sized bins: A cardinal (more precise) and an oblique (less precise) bin – both defined in allocentric coordinates. The cardinal bin included orientations from + 20º around 0º and + 20º around 90º, while the oblique bin included all the other orientations (+ 20º around both 45º and 135º). We calculated the circular standard deviation of recall errors across all trials in the oblique and cardinal bins for each participant and each head tilt condition. To calculate the *magnitude* of the oblique effect in each participant, we then subtracted oblique from cardinal bins. Because the bins are defined in allocentric coordinates, for head-upright trials the obliques have a larger circular SD and the cardinals have a smaller circular SD. This means that a *negative* cardinal-minus-oblique difference indicates a stronger oblique effect with the head upright. However, because the oblique effect tends to be anchored to the retinocentric reference frame, for head-tilted trials the obliques have a smaller circular SD (they are cardinal on the retinae) while the cardinals have a larger circular SD. This means that a *positive* cardinal-minus-oblique difference indicates a stronger oblique effect with the head tilted.

To quantify the repulsion effect at the level of individual participants, we divided the orientation space into two equally sized bins: A clockwise bin and a counterclockwise bin – again both defined in allocentric coordinates. The clockwise bin included orientations from 5º to 40º relative to each cardinal (so relative to 0º and to 90º), while the counterclockwise bins included orientations from -40º to -5º relative to each cardinal. Since no bias was expected for exact cardinal (0º and 90º) and obliques (45º and 135º), these orientations were excluded from this analysis. We calculated the signed circular mean of recall errors across all trials in the clockwise and counterclockwise bins for each participant and each head position condition, such that positive error reflect a clockwise bias relative to the correct response, and negative error a counterclockwise bias. To estimate the magnitude of the repulsion bias, we calculated the absolute difference in recall error between the clockwise and counterclockwise bins. Since the repulsion effect tends to stay anchored to an allocentric reference frame during both head-upright and head-tilted trials, a higher value always indicates a stronger repulsion bias for both conditions.

Finally, we wanted to explore how individual differences in neural decoding might be related to individual differences in oblique and repulsion effect strengths. To this end, we correlated each of these two behavioral effects (from the head-tilted trials) with our “relative decoding” metric (allocentric minus retinocentric, from the main cross-generalization analyses) at each timepoint during a trial, using a Spearman correlation coefficient.

#### Statistical significance testing

To determine statistical significance, we use two-tailed non-parametric permutation tests with 10000 iterations and an alpha level of 0.05, unless explicitly stated otherwise. To see if decoding accuracies were significantly above zero (i.e., when decoding orientation within the head-upright and head-tilted conditions), we used the sign-permutation test^67^. On each permutation, we randomly flipped the sign of each participant’s decoding accuracy with a 50% probability. These randomly flipped accuracies were used to calculate a one-sample *t-*statistic,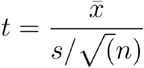, where and are the mean and standard deviation of decoding accuracies across participants, and is the sample size. This resulted in one ‘null’ decoding accuracy estimate per permutation, and a null-distribution of decoding accuracies across all permutations. The *t*-statistic of the true data was compared against the permuted null-distribution for each given timepoint and condition. To correct for multiple comparisons over time, we used a cluster-based correction. First, across all permutations for each timepoint, we found the t-value corresponding to the alpha-threshold. Based on these *t*-values for each timepoint, we found all significant timepoints in each permuted time course. For multiple adjacent significant timepoints, we calculated the sum of their *t*-values. The maximum summed *t*-value on each permutation was added to a null distribution of summed *t*-values. The *p*-value of each cluster in the true data was then calculated as the number of permuted summed *t*-values in the null distribution equal to or larger than the true summed *t*-value divided by the number of permutations. These cluster tests are run separately for the perceptual time period (decoding from voltage) and for the delay period (decoding from alpha).

For the cross-generalized decoding, the same approach was used for the relative decoding metric (allocentric minus retinocentric), where in this case a value significantly above zero would indicate an allocentric reference frame, while a value significantly below zero would indicate a retinocentric reference frame. For the relative decoding calculated from the early perception epoch, which looks at the internal consistency of individual participants’ reference frames, we used the same sign-permutation test, but instead of randomly flipping the sign of each participant’s average, we randomly flip the sign of each trial (for a given participant) to arrive at a within-subject *t*-statistic. For visualization of the circular mean, and to assess if it was closer to the allocentric or retinocentric frame, we computed 95% bootstrap confidence intervals with 10000 iterations by resampling participant-level circular means with replacement, and calculating the average circular mean on each iteration.

For the brain-behavior correlations, we ran permutation tests with 10000 iterations, randomly shuffling the behavioral effect strengths across participants on every permutation to break the participant-level brain-behavior relationship. To correct for multiple comparisons over time, we apply the same cluster-based correction as used for the decoding accuracy. To plot these correlations over time, we compute 95% confidence intervals using permuted correlation coefficients.

## Acknowledgements

Data acquisition was financed by NEI R01-EY025872 and NIMH R01-MH087214 to John T. Serences – whose unwavering and continued support is not to be underestimated, especially when we start shelving out grant monies to build rotating chin rests. CC is funded by grants for development of new faculty staff, the Rachadaphiseksomphot fund, and Chulalongkorn University. MVS, MJW, and RLR are funded by the Max Planck Society.

## Supplementary materials

**Supplementary Figure 1.**
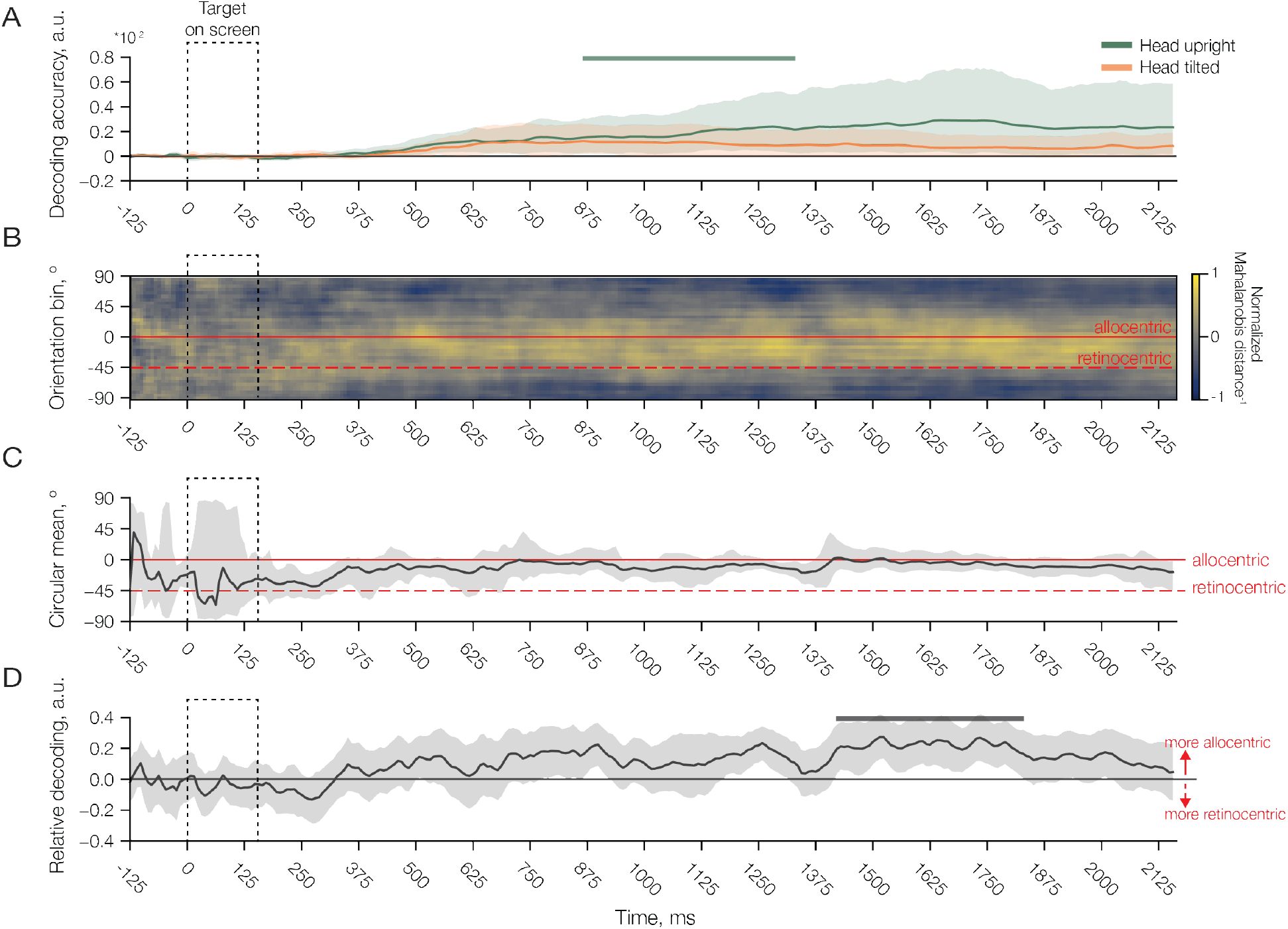
Decoding from the electrodes placed around the eyes (the EOGs). **A:** To assess the potential information contained in eye movements, we repeated the within-condition decoding analysis on the EOGs (upright in green, tilted in orange). We show a brief period of significant decoding when the head is upright, from 875-1300 ms after stimulus onset. There was no significant decoding in the head-tilted trials. This result suggests that eye movements did not meaningfully contribute to the EEG decoding results. **B:** Cross-generalized decoding heatmap. Despite there being little to no decoding within head tilt conditions (as in **A**), there does appear to be some clustering of smaller Mahalanobis distances (in yellow) later during the delay. **C:** Circular mean of the cross-generalized decoding, showing predominantly allocentric information in eye movements (though this is difficult to interpret in the absence of robust within-condition decoding). **D:** Decoding accuracy difference (allocentric minus retinocentric) for the cross-generalized decoding. Results are is in line with what we observe for the circular mean, in that they reveal a significant allocentric cluster later during the delay. This indicates that, if anything, the eyes move systematically with the orientation as it was shown on the screen. Error areas represent bootstrapped 95% confidence intervals of the mean, while solid bars/lines denoting significant decoding clusters (permutation test, n = 19, cluster-defining threshold *p* < .05, corrected significance level *p* < .05).

**Supplementary Figure 2.**
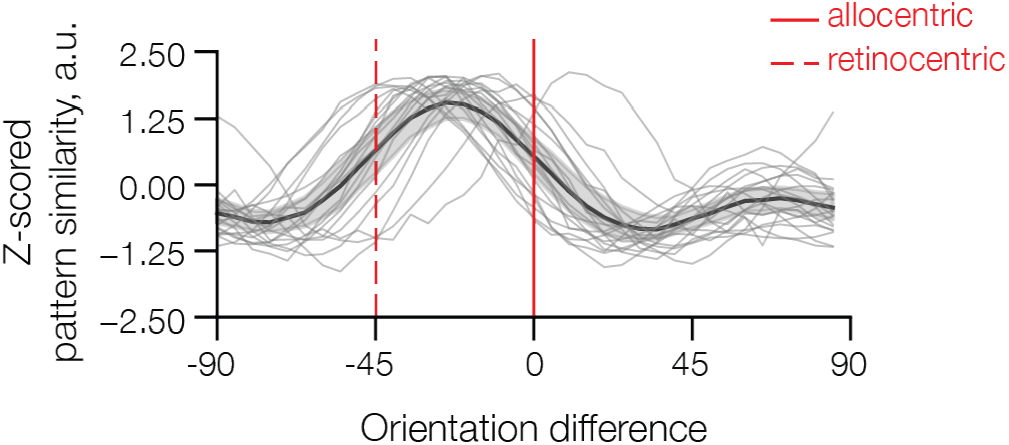
To probe the possibility that the reference frame was still retinocentric during the very earliest moments of perception (i.e., during the feedforward sweep), we applied a spatiotemporal decoder to the voltage signals from 50-100 after stimulus onset. This approach yields more robust decoding by incorporating temporal dynamics into a single decoding estimate. Shown here is the Mahalanobis curve during this early time period, with the group-average shown in dark grey (thicker line with error area), and individual participants shown in light gray (thinner lines). Dashed vertical lines reflect the target orientation in either the allocentric (solid red) and retinocentric (dashed red) reference frame. Note that while the target representation is in between both reference frames on average, there are strong inter-individual differences with the Mahalanobis curves of some participants clearly shifted towards a more allocentric or retinocentric reference frame.

**Supplementary Figure 3.**
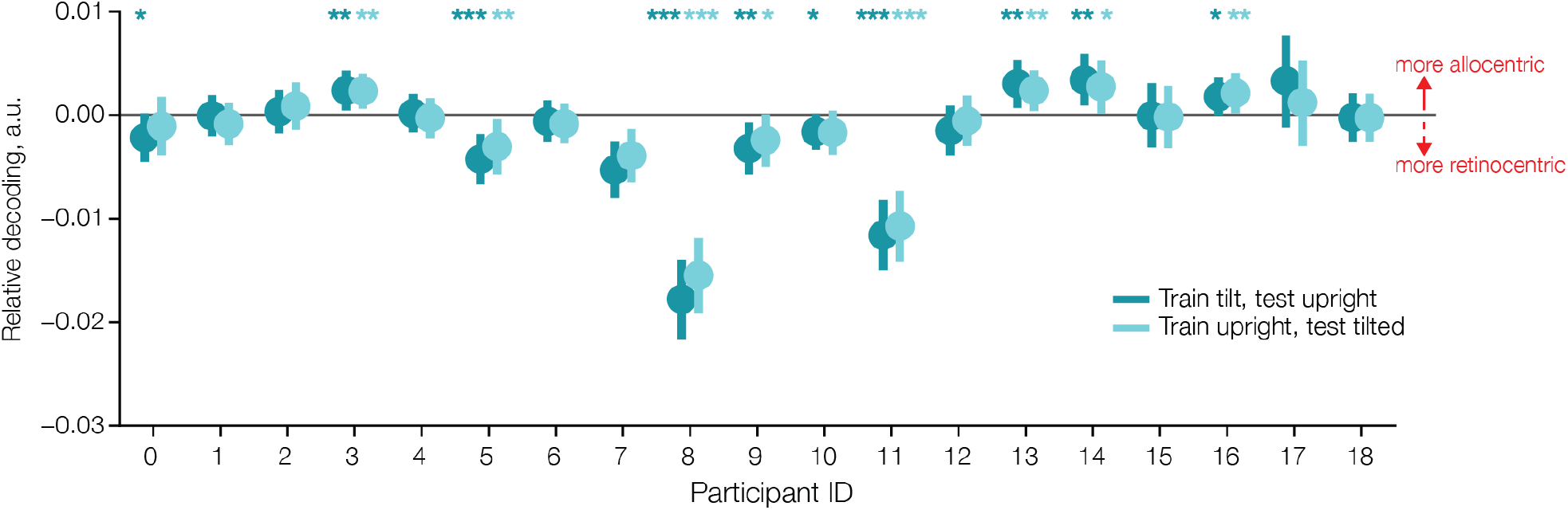
Individual participants’ mean relative decoding accuracies using the spatiotemporal decoder to the voltage signals from 50-100 after stimulus onset. For relative decoding, retinocentric decoding is subtracted from allocentric decoding, so that positive relative decoding denotes a more allocentric representation, while negative relative decoding denotes a more retinocentric representation. Here we show results for two cross-generalized decoders: one trained on upright trials and tested on tilted trials (light blue) and one trained on tilted trials and tested on upright trials (dark blue). Error bars indicate bootstrapped 95% confidence intervals, and asterisks indicate the significance level for each individual participant based on permutation testing (* for *p* < .05, ** for *p* < .01, and *** for *p* < .001). A participant was considered as allocentric if the mean relative decoding accuracies for both decoders were significantly above 0, retinocentric if they were significantly below 0, and “in between” otherwise.

